# Ascending vaginal infection in mice induces preterm birth and neonatal morbidity

**DOI:** 10.1101/2023.08.14.553220

**Authors:** Ashley K Boyle, Konstantina Tetorou, Natalie Suff, Laura Beecroft, Margherita Mazzaschi, Mariya Hristova, Simon N Waddington, Donald Peebles

## Abstract

Preterm birth (PTB; delivery <37 weeks), the main cause of neonatal death worldwide, can lead to adverse neurodevelopmental outcomes, as well as lung and gut pathology. PTB is commonly associated with ascending vaginal infection. Previously, we have shown that ascending *E. coli* infection in pregnant mice induces PTB and reduces pup survival. Here, we demonstrate that this model recapitulates the pathology observed in human preterm neonates, namely neuroinflammation, lung injury and gut inflammation. In neonatal brains, there is widespread cell death, microglial activation, astrogliosis and reduced neuronal density. We also validate the utility of this model by assessing efficacy of maternal cervical gene therapy with an adeno-associated viral vector containing human beta defensin 3; this improves pup survival and reduces *Tnfα* mRNA expression in perinatal pup brains exposed to *E. coli*. This model provides a unique opportunity to evaluate the therapeutic benefit of preterm labour interventions on perinatal pathology.

## Introduction

Preterm birth (PTB; delivery before 37 weeks) is the leading cause of neonatal morbidity and mortality worldwide (WHO, 1977; Liu *et al*., 2016). This is a serious global health problem, affecting approximately 11% of all pregnancies and 15 million babies a year (Chawanpaiboon *et al*., 2019; Unicef, 2017; Blencowe *et al*., 2012). Risk factors include extremes in maternal age, BMI, socioeconomic disadvantage, smoking, multiple pregnancies and a previous PTB (Boyle *et al*., 2017; Rubens *et al*., 2014).

Despite extensive research, treatment strategies have proved largely ineffective. This is likely due to the multifactorial nature of spontaneous PTB. The cervix, a physical and immunological barrier containing immune cells and producing antimicrobial peptides (AMPs), plays a key role in preventing bacterial invasion. However, at least 40% of spontaneous preterm births are associated with microbes, which are thought to ascend from the vagina into the uterus, via the cervix (Jones *et al*., 2009). The microorganisms commonly linked with PTB have relatively low pathogenicity and cases of chorioamnionitis are mostly polymicrobial. Species of *Mycoplasma*, *Ureaplasma*, *Fusobacterium* and *Streptococcus* have been found in the amniotic fluid in those most at risk (DiGiulio *et al*., 2010; DiGiulio, 2012). Vaginal *Lactobacillus iners* dominance has been linked to PTB risk in a UK population, while *Lactobacillus crispatus* dominance has been associated with term delivery (Kindinger *et al*., 2017; Bayar *et al*., 2020; Tabatabaei *et al*., 2019).

The fetal immune system is primed *in utero* through exposure of the placenta, lungs, gut and skin to microbes, as well as the vertical transmission of maternal immune cells (Mishra *et al*., 2021; Stelzer *et al*., 2021). When delivered, babies exposed to infection/inflammation have an increased risk of periventricular leukomalacia, intraventricular haemorrhage and early-onset neonatal sepsis. These conditions increase the likelihood of long term complications resulting in neurodevelopmental disorders, such as cerebral palsy, and respiratory related conditions, such as bronchopulmonary dysplasia (BPD), and necrotising enterocolitis (NEC) (Jung *et al*., 2020; Humberg *et al*., 2020).

Previously, we demonstrated that PTB can be induced in mice by ascending vaginal infection with a pathogenic *Escherichia coli* (*E. coli*) strain associated with human neonatal meningitis. We showed that the proportion of pups born alive was reduced and in surviving pups there was evidence of neuroinflammation (Suff *et al*., 2018). We used this model to investigate a novel therapeutic, demonstrating that adeno-associated viral (AAV) vector gene therapy to increase levels of the AMP human beta defensin 3 (HBD3) in the cervix can reduce ascending infection and improve pup survival (Suff *et al*., 2020).

Here, we characterise the fetal and neonatal outcomes following exposure to ascending vaginal infection, assessing the fetal lungs and gut, as well as further interrogating fetal and neonatal neuropathology. We also preventatively treat dams with cervical HBD3 gene therapy and assess the impact of this on perinatal neuroinflammation.

## Material and methods

### AAV vector production

An AAV8 bicistronic vector was produced encapsidating a single-stranded DNA sequence containing the HBD3 gene followed by the engineered green fluorescent protein (eGFP) gene, each under the transcriptional activity of a separate cytomegalovirus (CMV) promotor. An AAV8 vector containing CMV-eGFP was used as a vehicle control.

AAV production and purification followed standard procedures. Briefly, HEK293T cells were transfected with pAAV-CMV-hDEFB103B-CMV-eGFP or pAAV-CMV-eGFP (Vector Biolabs, Malvern, USA), using polyethylenimine (PEImax, Polysciences Inc, Warrington, PA, USA). Cells were incubated at 37°C for 72 hours then collected, centrifuged and freeze-thawed in lysis buffer. Both cell supernatant and cell lysate were benzonase-treated then clarified via centrifugation and 0.22μm filtration prior to purification. AAVs were purified by high-performance liquid chromatography (HPLC; ÄKTAprime plus, GE Healthcare, Buckinghamshire, UK) with a POROS™ CaptureSelect™ AAV Resin (Thermo Fisher Scientific, Oxford, UK) column. Purified vector fractions were dialysed against phosphate-buffered saline (PBS) overnight (Slide-A-lyzer dialysis cassette 10,000 MWCO, Thermo Fisher Scientific). Viral genome titres were quantified by quantitative RT-PCR (qRT-PCR).

### Mouse model of ascending vaginal infection

All animal studies were conducted under a UK Home Office License (PAD4E6357) in line with the Animal Scientific Procedures Act (1986) and following the ARRIVE guidelines. Temperature (19– 23°C) and humidity (∼55%) were tightly controlled at all times, with constant 12 hour light/dark cycles. Virgin female C57BL/6 Tyr^c-2J^ mice (Charles River, Kent, UK) aged 6-12 weeks were time mated (embryonic day (E) 0.5 designated when vaginal plug was found).

On E16.5, mice were anaesthetised by the inhalation of isoflurane (5% for induction, 1.5% for maintenance in oxygen). *E. coli* K1 A192PP-*lux2*, modified to contain the lux operon from *P. luminescens* (Cronin *et al*., 2012), was delivered intravaginally to induce preterm delivery as previously described (Suff et al 2018). Briefly, 20µL of midlogarithmic-phase *E. coli* (1 x 10^2^ CFU resuspended in PBS) or PBS (vehicle control) was delivered into the vagina of pregnant mice using a 200mL pipette tip, followed by 20µL of 20% Pluronic® F-127 gel (Sigma-Aldrich, Dorset, UK) to prevent leakage. Animals were allowed to recover from anaesthesia then caged individually and monitored by CCTV for signs of labour and delivery of pups. Time to delivery was recorded as the number of hours from the time of intravaginal administration of *E. coli* or PBS to the delivery of the first pup. The number of live/dead pups was recorded within 24 hours of their delivery and a percentage calculated per litter.

In a separate cohort, pregnant C57BL/6 Tyr^c-2J^ females (6-12 weeks) were anesthetised on E13.5 and 10µL of AAV8-HBD3 or AAV8-GFP was delivered intravaginally using a sterile 200µL pipette tip (1 × 10^12^ genomic copies/mL diluted in PBS) followed by 10µL of AK012 thermosensitive gel (PolySciTech, Indiana, USA). Administration of *E. coli* or PBS on E16.5 was then performed as already described.

### Bioluminescent imaging

Mice were imaged for 2 minutes using a cooled charged-coupled device camera (IVIS Lumina; Perkin Elmer, Coventry, UK). Dams were anaesthetised with isoflurane as above.

### Tissue collection and processing

Perinatal pups were sacrificed by decapitation on E18.5 (48 hours after intravaginal infection) to harvest brain, lungs and gut (1-3 pups per litter collected from the right horn). Dams and postnatal day (P) 7 and P14 pups were anaesthetised using isoflurane, the right atrium was incised and PBS was injected into the left ventricle. P7 brains were cut into three coronal sections: the forebrain, the mid brain region and the hindbrain/cerebellum. Tissues for qRT-PCR and ELISA analyses were snap-frozen and stored at −80°C until required. For immunohistochemistry, brain and lung samples were fixed in 4% paraformaldehyde for 24-48 hours then stored in 30% sucrose at 4°C. Tissues were frozen on dry ice and a cryostat (LEICA CM1900, Newcastle upon Tyne, UK) was used at −20°C to cut 50 serial 40µm sections of the tissues, beginning from the fusion of the corpus callosum of the brain. Frozen sections were stored at −80°C.

### RNA extraction, cDNA synthesis and qRT-PCR

Total RNA was extracted from E18.5 brains, lungs and gut and P7 forebrain and mid brain samples by RNAeasy mini kit (Qiagen, Germantown, MD, USA) according to the manufacturer’s instructions, and quantified on a FLUOstar Omega Microplate Reader (BMG Labtech Ltd, Aylesbury, UK). RNA (400 ng) was reverse transcribed by using the High Capacity cDNA Reverse Transcription Kit (Thermo Fisher Scientific). For qRT-PCR, primers for *Gapdh* (Forward: ACGGCAAATTCAACGGCAC, Reverse: TAGTGGGGTCTCGCTCCTGG) and *Il-1β* (Forward: AACCTGCTGGTGTGTGACGTTC, Reverse: CAGCACGAGGCTTTTTTGTTGT) were designed and purchased from Integrated DNA Technologies (Leuven, Belgium) or predesigned TaqMan gene expression assays were used (Thermo Fisher Scientific, Table 1). All qRT-PCR analyses were performed on a QuantStudio 6 Flex Real-Time PCR System (Applied Biosystems, Thermo Fisher Scientific). Target mRNA expression was normalised for RNA loading using *Gapdh* (Integrated DNA Technologies), and the mRNA concentration in each sample was calculated relative to the vehicle control group (PBS or AAV8-GFP + PBS) using the 2^-ΔΔCt^ method of analysis.

**Table 1.**
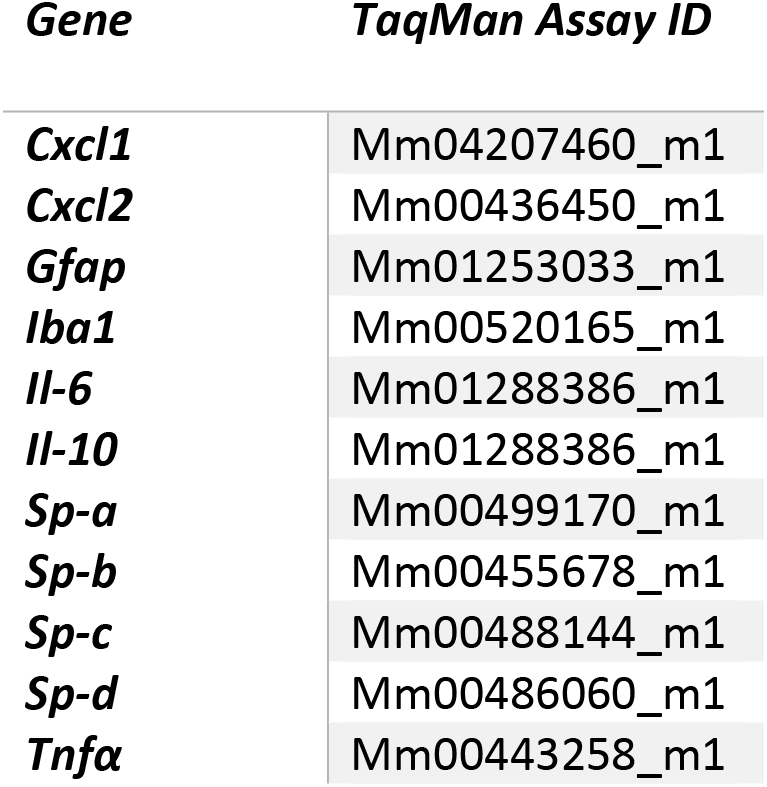
Predesigned TaqMan gene expression assays.

| *Gene* | *TaqMan Assay ID* |
| --- | --- |
| ***Cxcl1*** | Mm04207460_m1 |
| ***Cxcl2*** | Mm00436450_m1 |
| ***Gfap*** | Mm01253033_m1 |
| ***Iba1*** | Mm00520165_m1 |
| ***Il-6*** | Mm01288386_m1 |
| ***Il-10*** | Mm01288386_m1 |
| ***Sp-a*** | Mm00499170_m1 |
| ***Sp-b*** | Mm00455678_m1 |
| ***Sp-c*** | Mm00488144_m1 |
| ***Sp-d*** | Mm00486060_m1 |
| ***Tnfα*** | Mm00443258_m1 |

### Enzyme-linked immunosorbent assay

E18.5 brains were homogenised in lysis buffer (RIPA buffer; Millipore, Sigma-Aldrich) containing 1 Complete Protease Inhibitor Cocktail Tablet (Roche, Basel, Switzerland) using a Tissue Lyser II (Qiagen) at 30 Hz. The samples were incubated on ice for 5 minutes and then centrifuged at 10,000 g for 10 minutes at 4°C. The lysates were stored at −80°C. Total protein was quantified by using the DC protein assay (Bio-Rad, Hercules, CA, USA) according to the manufacturer’s instructions. The V-PLEX Proinflammatory Panel 1 Mouse Kit (Meso Scale Diagnostics, Maryland, USA) was used to quantify inflammatory mediator secretion according to the manufacturer’s guidelines and measured on a Meso Sector S 600mm. For E18.5 lungs and gut, total protein quantification was performed using the Pierce BCA Protein Assay kit (Thermo Fisher Scientific, Glasgow, UK), then the MILLIPLEX mouse cytokine/chemokine magnetic bead panel (Luminex® Multiplex Assay, R&D Biosystems) was used according to the manufacturer’s guidelines and measured on a Luminex® instrument with xPONENT software.

### Histology and Immunohistochemistry

Histological staining included terminal transferase mediated d-UTP nick end labelling (TUNEL) with Co/Ni enhancement (Roche, UK) performed following manufacturer’s instructions, cresyl-violet (Nissl) staining, and hematoxylin and eosin staining (H&E). For brain histology and immunohistochemistry, five sections per brain (400µm apart) were rehydrated in distilled H_2_O and left to dry. Immunohistochemical staining was performed as previously described (Hristova *et al*., 2010). Briefly, tissue sections were incubated overnight with the following primary antibodies: rabbit polyclonal anti-glial fibrillary acidic protein (GFAP) (1:6,000, DAKO, UK), rabbit polyclonal anti-allograft inflammatory factor 1 (IBA1) (microglia, 1:2,000, Wako, Japan), rabbit anti-myelin basic protein (MBP) (1:200, Abcam, Cambridge, UK), or mouse monoclonal anti-NeuN (neurons, 1:15,000, Millipore, UK). The following day, sections were incubated with either biotinylated goat anti-rabbit or anti-rat (1:100, Vector Laboratories, UK) secondary antibodies for 1 hour, then incubated with Avidin-Biotinylated horseradish peroxidase complex (Vector Laboratories) followed by diaminobenzidine (DAB)/ H_2_O_2_ (Fisher Scientific).

Double immunofluorescence for activated pro- and anti-inflammatory microglial cells was performed as above with rat anti-CD86 (M1; 1:1600 dilution, Dako) and goat anti-CD206 (M2; 1:400 dilution, Dako) primary antibodies followed by biotinylated donkey anti-rat (1:200 dilution, Vector Laboratories) and donkey anti-goat-488 (1:200 dilution, Vector Laboratories,) secondary antibodies, respectively. Sections were washed 3 times in 10mM phosphate buffer (PB), then incubated with Texas Red (1:1000, Bio-Techne, Abingdon, UK) and rabbit anti-goat-488 (1:200) respectively. Coverslips were mounted with medium containing DAPI (Vector Laboratories).

For lung analyses, two or three sections of lung per fetus or neonate (400µm apart) were rehydrated in distilled H_2_O then stained with Nissl or immunohistochemical labelling was performed as above with anti-mouse Ly6G primary antibody (BioLegend, London, UK) and goat anti-rabbit secondary antibody.

### Histological assessments

Assessments were performed on the following regions: cortex, pyriform cortex, external capsule, striatum, hippocampus and thalamus. The corpus callosum was analysed for E18.5 brains in addition to or in place of the external capsule.

#### Optical luminosity

Optical luminosity values (OLV) were generated to measure reactive astrogliosis and myelination through GFAP and MBP immunoreactivity, respectively. All regions were assessed for GFAP and the striatum and external capsule were assessed for MBP. Images were captured in three optical fields using a Sony AVT-Horn 184 3CCD camera (20x magnification, 24bit RGB, 760 x 570 pixel resolution) then imported into Optimas 6.51 (Media Cybernetics Inc., MD, USA) to determine the mean and standard deviation (SD) of the OLV. To obtain the intensity of the staining for each image, the SD was subtracted from the mean OLV and the resulting values were normalised by subtracting the OLV of the glass surrounding the tissue section (Möller *et al*., 1996).

#### Ramification index

IBA1 positive microglial cell bodies and branch density were assessed using a 0.049 mm × 0.049 mm square grid (40x magnification) placed in three fields for all brain regions. The number of cell bodies within the grid were counted (C) and the average number of branches crossing the three horizontal and three vertical 0.49 mm gridlines (B). The microglial ramification index was calculated as (B2/C).

#### Stereology for brain and lung tissues

TUNEL (20x magnification), NeuN (40x magnification) and Ly6G positive cells (40x magnification) were manually counted using ImageJ. Brain width, cortex thickness and hippocampal area and lung alveoli area were measured using the free-hand tool (40x magnification). Three optical fields were assessed per brain region/lung tissue section and an average found.

CD86 and CD206 immunoreactivity were quantified in three fields per region of interest using a fluorescent microscope (40x magnification; Leica). Positive staining and co-localised staining were counted using CellProfiler with a MATLAB designated code, then verified using the ImageJ multi-point tool. An average was found per region, per tissue.

### Statistics

Data were analysed by using GraphPad Prism v.8 (GraphPad Software, La Jolla, CA, USA). Time to delivery was analysed by an unpaired t test with Welsh’s correction or one-way ANOVA with Sidak’s multiple comparisons test. The percentage data for live-born pups were analysed by performing an arcsine transformation on the proportions, to normalise the binomial distribution, followed by an unpaired t test with Welsh’s correction or one-way ANOVA with Sidak’s multiple comparisons test. Pup survival was analysed by Log-rank (Mantel-Cox) test. qRT-PCR data were then analysed by using an unpaired t test with Welsh’s correction or one-way ANOVA with Tukey’s multiple comparisons test. All ELISA data were log transformed followed by unpaired t tests with Welsh’s correction. For histology and immunohistochemistry, either a Mann-Whitney test or multiple t tests with Welsh’s correction were used. Scoring for histology and immunohistochemistry was blinded where possible. Values of p<0.05 were considered statistically significant.

## Results

### Ascending vaginal infection with *E. coli* reduces gestational length as well as the percentage of pups born alive and pup survival over 7 days

On E16.5, 1 x 10^2^ CFU of bioluminescent *E. coli* or 20µL PBS control was administered to dams intravaginally. Within 24 hours, imaging shows the ascension of the bacteria from the vagina into the uterus (Figure 1A). Gestational length was significantly reduced when dams received *E. coli* compared to PBS-treated mice (Figure 1B; mean time to delivery 40.3 ± 7 hours versus 52.7 ± 12.9 hours for *E. coli* and PBS group, respectively; p=0.018). The average percentage of pups born alive was reduced by almost 30% following *E. coli* exposure (71.4 ± 34% vs 100 ± 0% in the PBS group; p=0.0054) and postnatal survival over days 7 was also reduced (p<0.0001).

**Figure 1.**
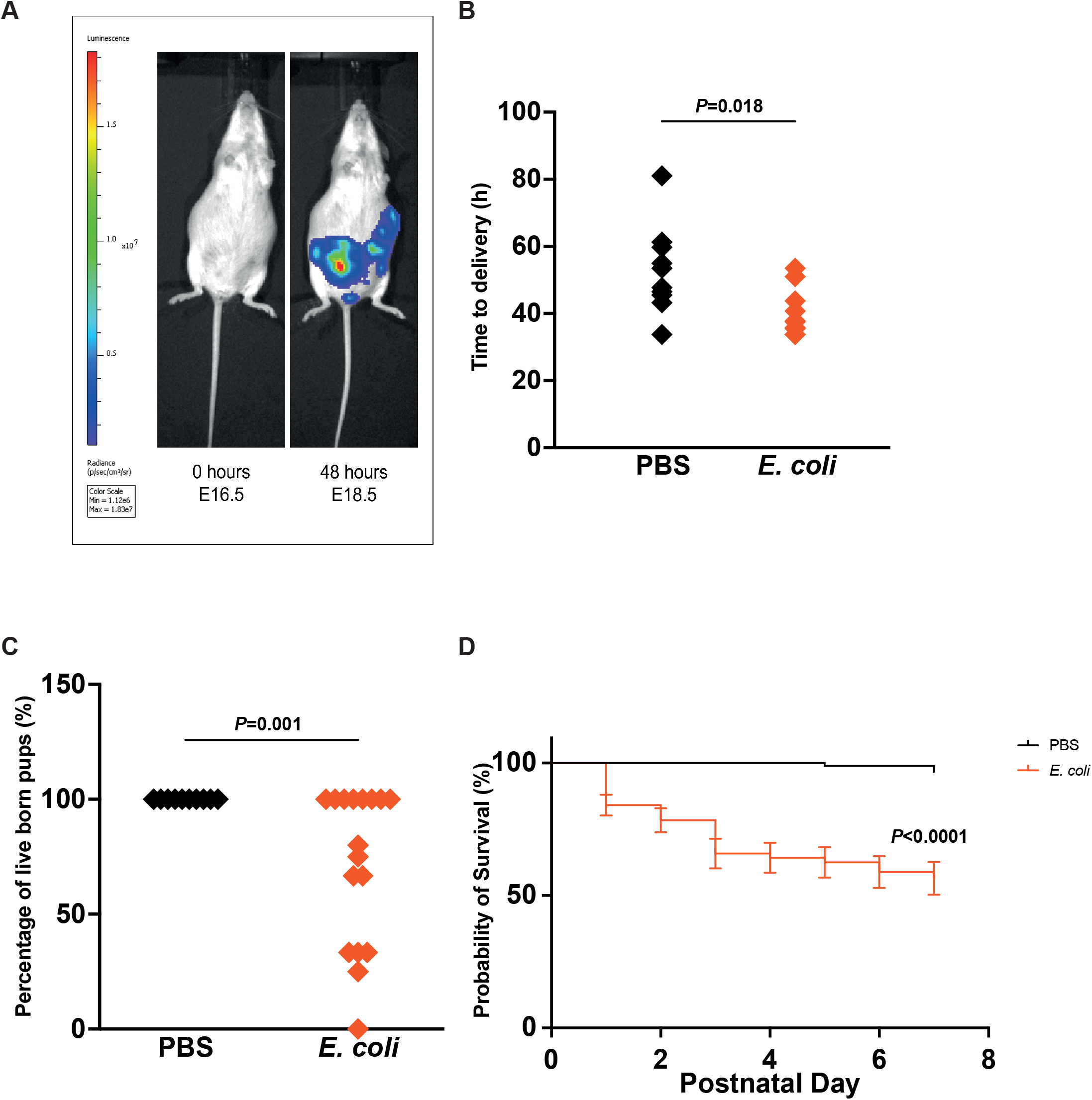
*E. coli* ascends from the vagina into uterus to induce PTB and reduce pup survival. After 48 hours, bioluminescent *E. coli* is visable in the uterus (A). *E. coli* reduces the time to delivery (B), the percentage of live born pups (C) and the long term survival of pups (D).

### Exposure to ascending vaginal infection *in utero* induces perinatal neuroinflammation

On E18.5, 48 hours after maternal vaginal infection with *E. coli*, perinatal pup tissues were collected and assessed for inflammation and pathology. Inflammatory mediator mRNA expression was upregulated with *E. coli* exposure (Figure 2A); *Tnfα* (p=0.025), *Il-1β* (p=0.01) and *Il-6* (p=0.034). TNFα secretion was also significantly elevated (Figure 2B; 0.6 ± 0.2 pg/mL vs 4.9 ± 12.3 pg/mL in PBS and *E. coli* pups, respectively; p=0.003). The secretion of IL1-β, IL-2, IL-4, IL-5, IL-6, IL-10, CXCL1 and IFNγ were unchanged (Figure 2B, Supplementary figure 1A).

**Figure 2.**
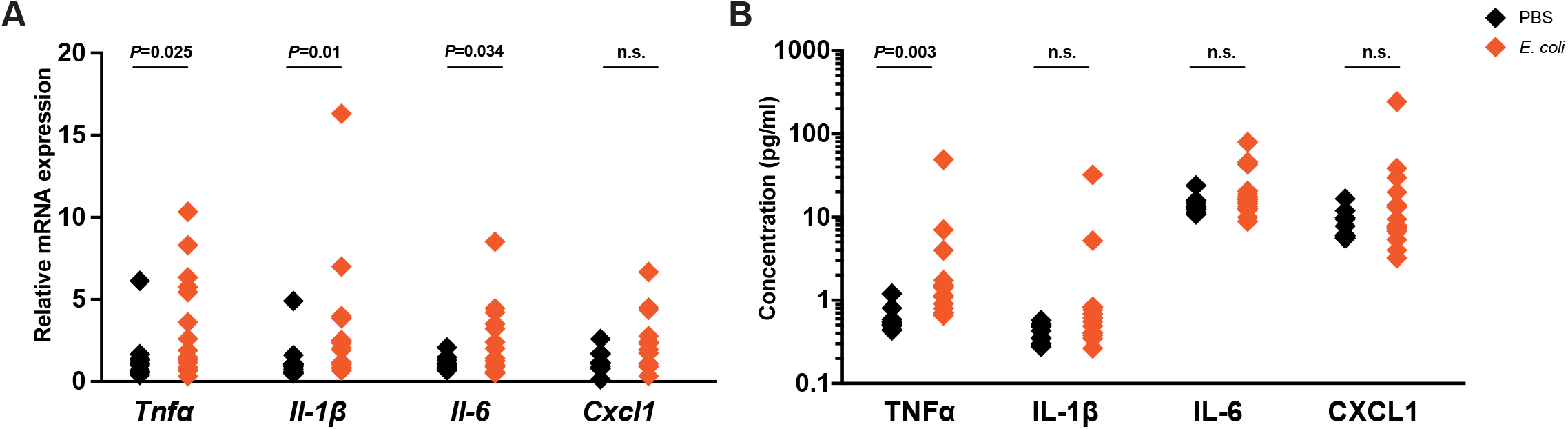
Ascending *E. coli* infection leads to perinatal neuroinflammation 48 hours after exposure. Proinflammatory mediators are upregulated at the gene (A) and protein (B) level in the perinatal pup brain following maternal *E. coli* infection.

Histological analyses suggested a trend towards TUNEL+ cell death, particularly in the corpus callosum and striatum (Supplementary figure 1B) and trends for reduced microglial ramification were seen in the cortex and thalamus (Supplementary figure 1C). The average numbers of NeuN+ neurons in the cortex and hippocampus were not altered by *E. coli* exposure at this time point and we did not observe any morphological changes (Supplementary figure 1D-G). There was a reduction in the average number of CD86+ cells (M1 macrophages) in the thalamus of the brain but there were no alterations in other brain regions, nor in CD206+ staining (M2 macrophages) or CD86/CD206 co-localisation (Supplementary figure 2).

### *E. coli* exposure *in utero* leads to neonatal neuropathology

At P7, *Iba1* mRNA, a marker of microglia, was upregulated in the forebrain region (p=0.016), as was *Il-1β* mRNA (p=0.001) (Figure 3A). In the mid brain region, mRNA expression was (Figure 3B); *Tnfα* (p<0.0001), *Il-1β* (p<0.0001), *Il-6* (p=0.028), *Cxcl1* (p=0.005), *Iba1* (p<0.0001) and *Gfap* (p=0.0006).

**Figure 3.**
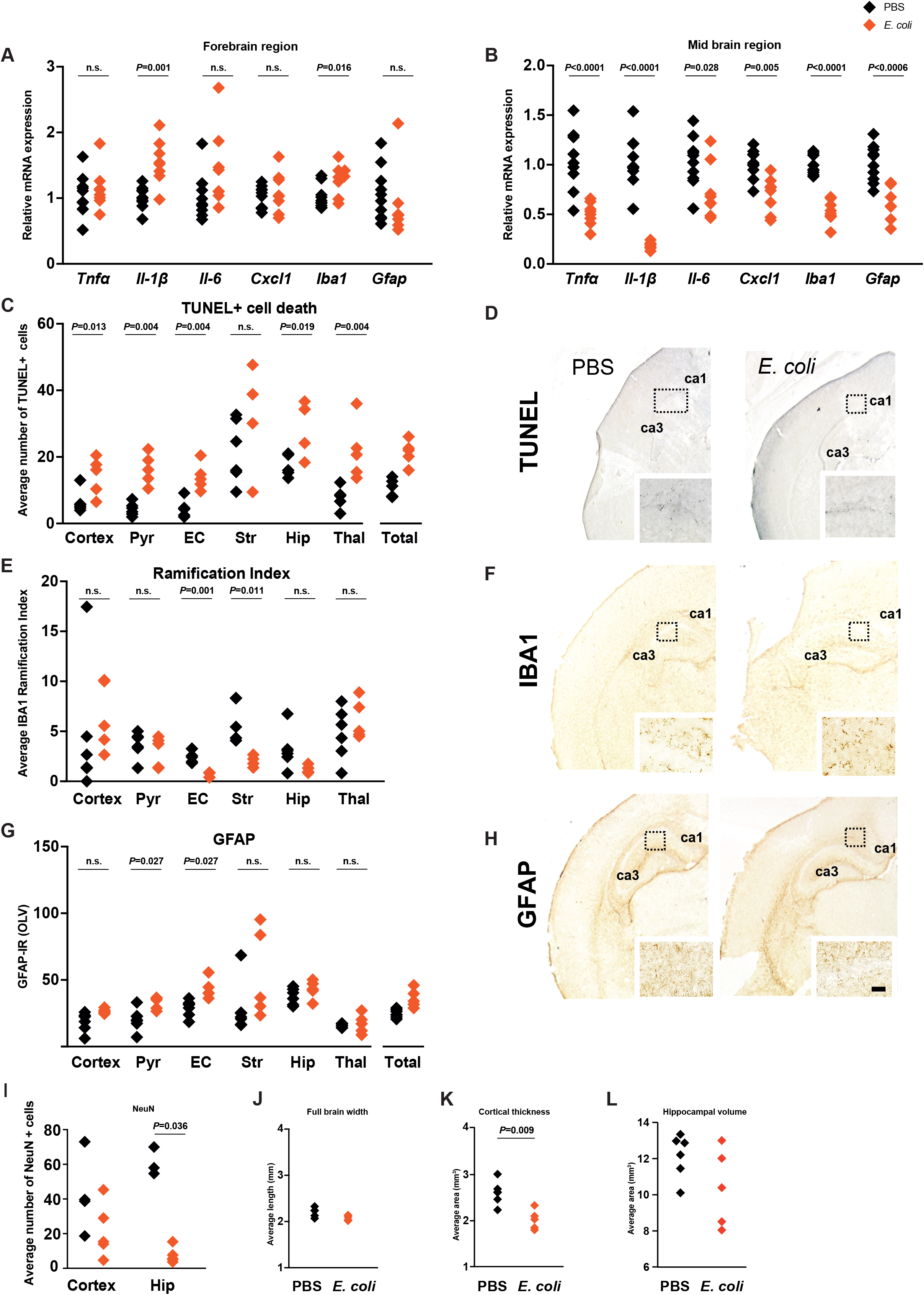
Neuropathology is evident in neonates exposed to *E. coli in utero*. Inflammatory mediators *Il-1β* and *Iba1* are upregulated in the forebrain (A), while all inflammatory mediators are downregulated in the mid brain region (B). The average number of TUNEL+ cells is increased in most brain regions (C) and IBA1+ cell ramification is reduced in the external capsule and striatum of the brain in the *E. coli* group (E). GFAP is increased in the pyriform cortex and the external capsule (G). The average number of NeuN+ cells is reduced the hippocampus (I) and the cortical thickness is reduced in the *E. coli* group (K). Pyriform cortex (Pyr), external capsule (EC), striatum (Str), hippocampus (Hip), thalamus (thal), total (all regions), cornu ammonis (CA), optical luminosity values (OLV). Scale bar = 59.5µm.

Histological analyses found widespread TUNEL+ cell death in *E. coli* pup brains: cortex (p=0.013), pyriform cortex (p=0.004), external capsule (p=0.004), hippocampus (p=0.019) and thalamus (p=0.004) (Figure 3C-D). We observed a reduction in microglial ramification (Figure 3E-F) in the external capsule (p=0.001) and striatum (p=0.011). GFAP, a marker of astrogliosis, was significantly increased in the pyriform cortex (Figure 3G-H; p=0.027) and external capsule (p=0.027). *E. coli*-exposed pups also had reduced numbers of NeuN+ neurons in the hippocampal region of the brain (Figure 3I; p=0.036) and cortical area was reduced in these pups (Figure 3K, p=0.009).

At P14, neuronal density was reduced in the cortex (Figure 4A; p=0.032) and the hippocampus (p=0.016). There was no longer evidence of TUNEL+ cell death or astrogliosis and MBP staining intensity was not significantly altered (Figure 4B and supplementary figure 3).

**Figure 4.**
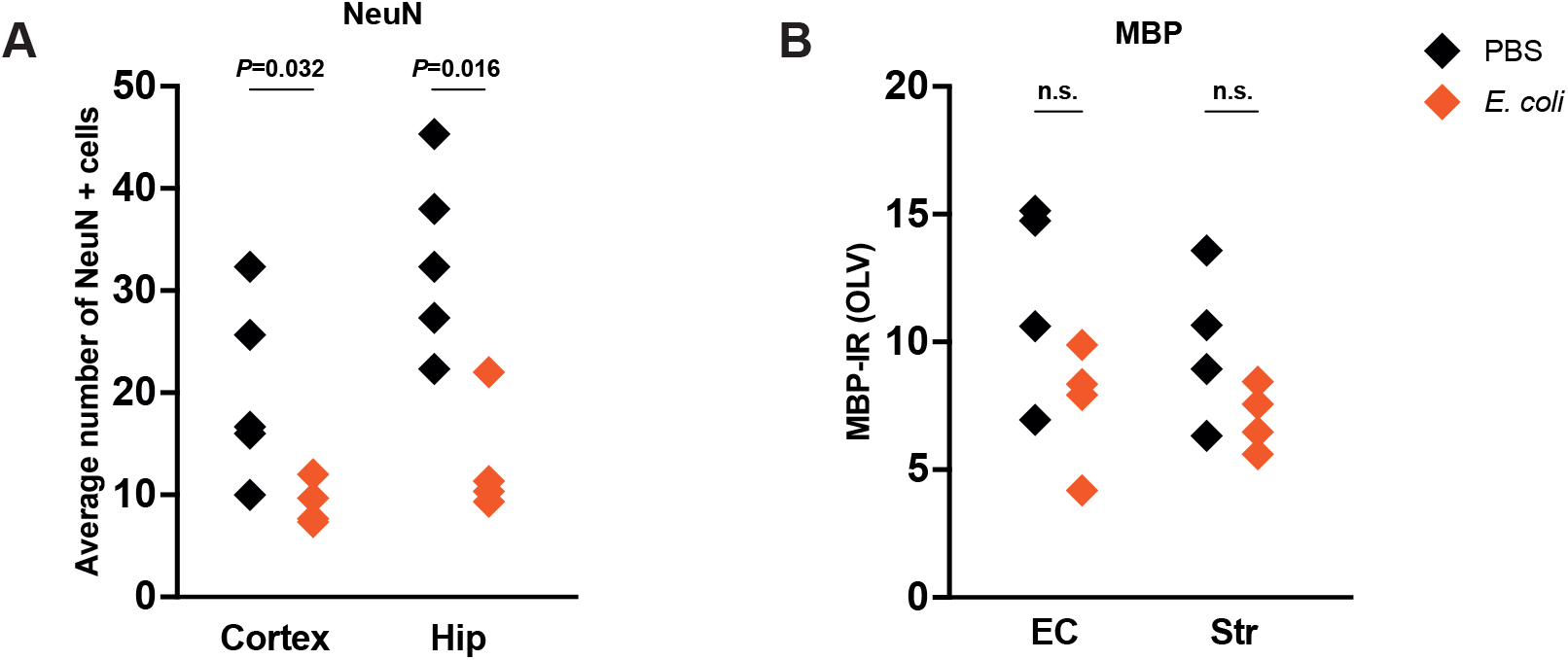
Neuronal density is reduced at P14 following *in utero E. coli* exposure. The average number of NeuN+ cells is reduced in cortex and hippocampus (A). There is no significant change to MBP luminosity (B). External capsule (EC), striatum (Str), hippocampus (Hip), optical luminosity values (OLV).

### Exposure to ascending vaginal infection *in utero* induces inflammation in the perinatal lungs and gut

#### Lung

Surfactant protein mRNA expression was downregulated in the lungs of pups from *E. coli*-treated dams (Figure 5A; *Sp-a* (p=0.023), *Sp-b* (p=0.034), *Sp-c* (p=0.006)). We also observed pup-dependent elevation of inflammatory mediator mRNA expression but, overall, there was no significant change to any of these mRNA markers compared to the PBS group (Figure 5B). However, IL-6 (p=0.035), CXCL1 (p=0.02) and Monocyte Chemoattractant Protein 1 (MCP1; p=0.004) were increased in these mice (Figure 5C). Additionally, we observed a significant increase in Ly6G-positive neutrophil infiltration in the lungs of *E. coli*-exposed pups (Figure 5D-E; p=0.011) and the mean alveolar area was increased (Figure 5F-G; p=0.003). Neutrophil influx and morphological changes were no longer evident at P7 (Supplementary figure 4).

**Figure 5.**
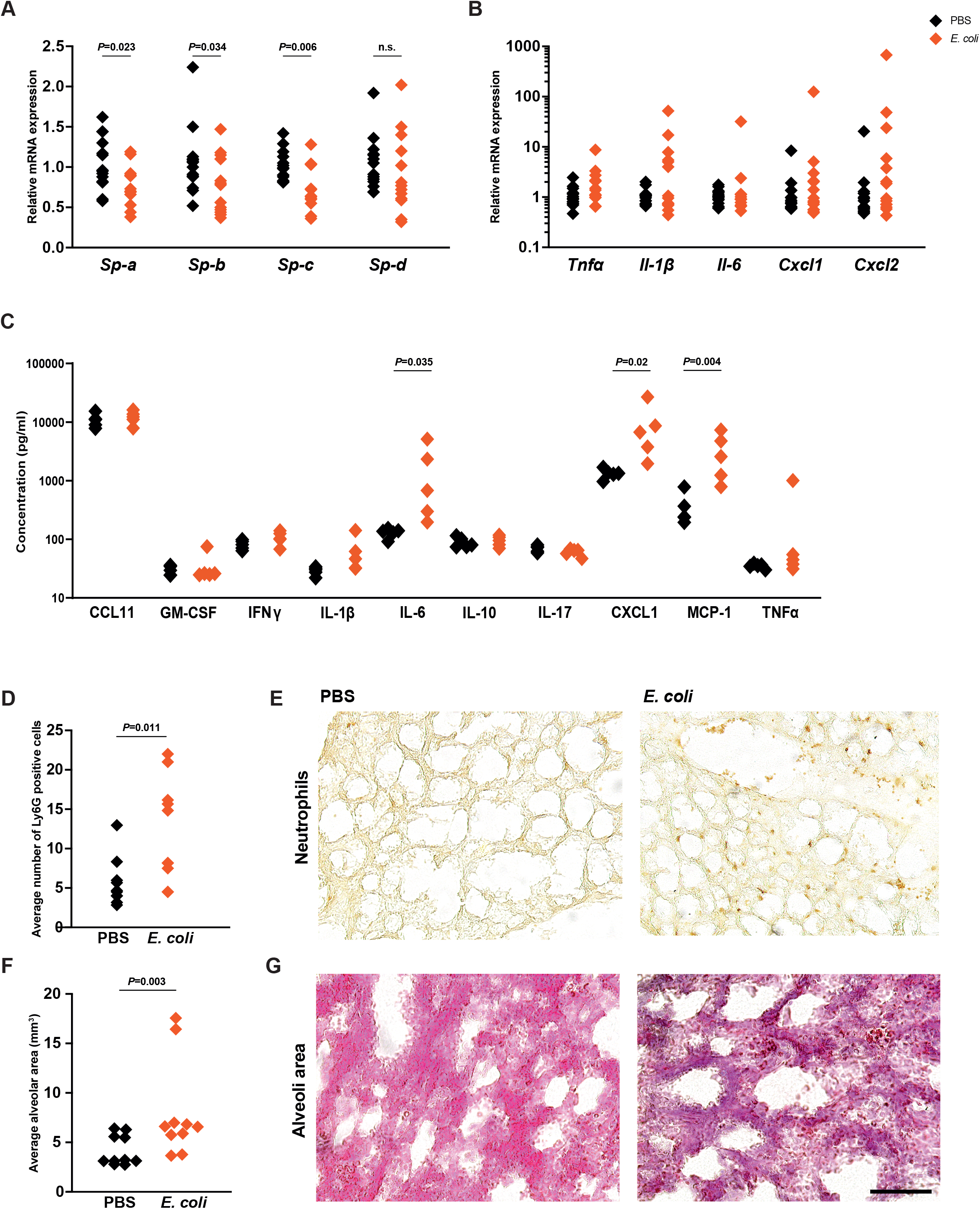
There is evidence of injury to the perinatal lungs following *E. coli* exposure. Surfactant protein mRNA expression is reduced in the *E. coli* group (A). A trend for increased inflammatory mediator expression can be seen at the mRNA level (B) and significant increases are observed at the protein level (C). The average number of Ly6G+ cells is increased (D-E) and the average alveolar area is increased in the *E. coli* group (F-G). Scale bar = 59.5µm.

#### Gut

In the gut, *Tnfα* (p=0.007), *Cxcl1* (p=0.031) and *Cxcl2* (p=0.038) mRNA expression were elevated but there was no impact on *Il-1β*, *Il-6* or *Il-10* mRNA expression or on the secretion of inflammatory proteins (Figure 6A-B).

**Figure 6.**
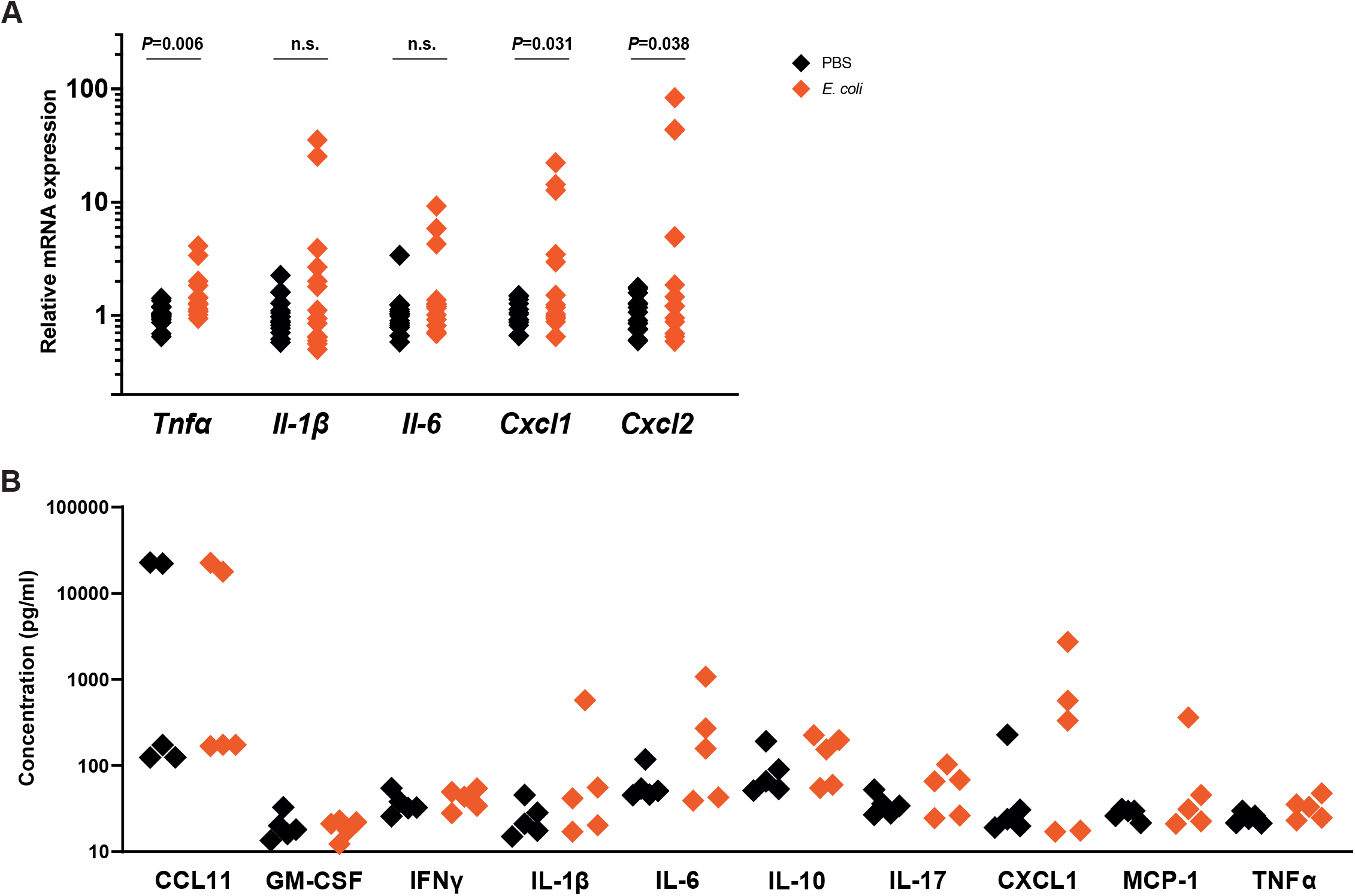
*E. coli* induces perinatal gut inflammation. Inflammatory mediator mRNA expression is increased in the gut (A). There is no change to the protein expression of these mediators (B).

### Maternal AAV8-HBD3 cervical gene therapy improves pup survival but does not increase gestational length

To validate the utility of the model, we administered AAV8-HBD3. Firstly, we evaluated the effect on PTB. The gestational length of infected dams was not increased with maternal AAV8-HBD3 treatment compared to the AAV8-GFP + *E. coli* group (42.54 ± 6 hours versus 46.55 ± 5 hours, respectively; Figure 7A). The percentage of live born pups was significantly increased in comparison to the AAV8-GFP + *E. coli* group (Figure 7B; 85.2% vs 58.6%, respectively; p=0.043) and pup survival was improved over 7 days (Figure 7C; p=0.002). This is consistent with our previous data using this treatment (Suff *et al*., 2020).

**Figure 7.**
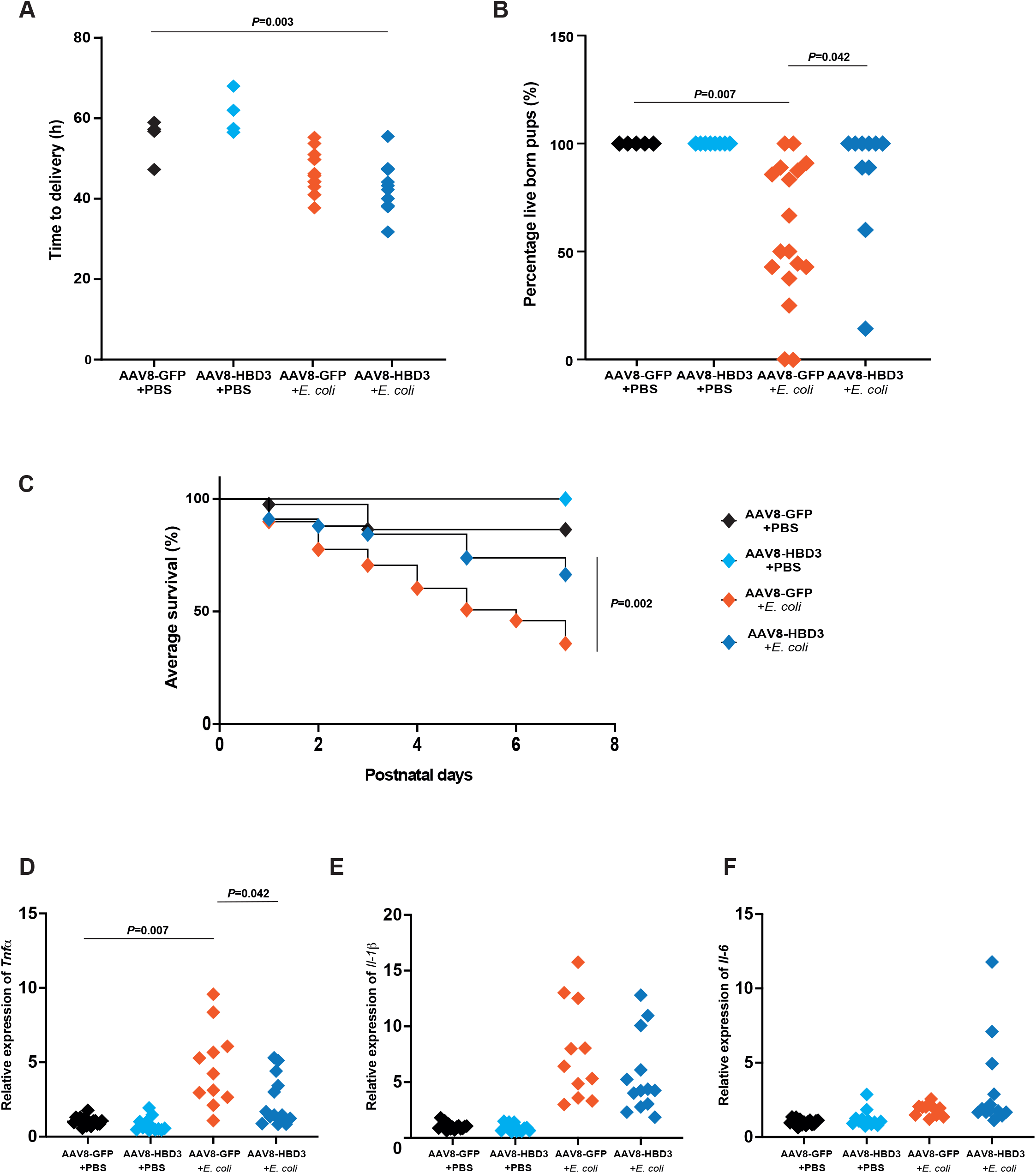
Cervical HBD3 gene delivery improves pup survival and reduces *Tnfα* in the perinatal brain. HBD3 treatment does not impact the length of gestation (A) but it increases the percentage of live born pups (B) and long term pup survival (C). *Tnfα*, but not *Il-1β* and *Il-6*, is reduced in the perinatal brain following maternal cervical HBD3 gene delivery (D-F).

This model also allowed us to examine the therapeutic effect on perinatal brain pathology. AAV8-HBD3 reduced *Tnfα* mRNA expression in the E18.5 pup brains exposed to *E. coli* compared to control vector (p=0.008) but did not significantly alter *Il-1β* or *Il-6* mRNA expression (Figure 7D).

## Discussion

PTB, and the resulting neonatal morbidity and mortality, is a serious global health issue for which preventative treatments are largely ineffective. It is essential to establish clinically relevant models to recapitulate not only preterm delivery but also the impact this has on the developing fetus in order to develop innovative interventions. As we have previously reported, it is possible to replicate ascending vaginal infection, the most common cause of PTB, in mice. Here, we demonstrate that we are also modelling neuropathology, lung injury and gut inflammation: key neonatal morbidities associated with prematurity. Furthermore, we used this model to investigate cervical gene delivery of HBD3, revealing that this intervention can significantly increase pup survival while reducing *Tnfα* expression in the perinatal brain.

### Modelling ascending vaginal infection and PTB

Our mouse model is the only one to our knowledge that recapitulates the human scenario whereby PTB is induced by live bacterial ascending infection, which also reduces pup survival and causes neonatal morbidity in the survivors. Mice are commonly utilised to model human PTB. There are a number of methods used to induce labour but by far the most common method is via lipopolysaccharide (LPS) administration (McCarthy *et al*., 2018). However, there is huge heterogeneity among models, which differ in the mouse strain used, LPS serotype, LPS dose, mode of delivery and day of induction (Miller *et al*., 2022). LPS models have several disadvantages. The use of a bacterial product rather than live bacteria means it is a model of inflammation not infection and the clinical scenario of ascension through the cervix cannot be recapitulated due to LPS immotility. In addition, the dose required to induce PTB is often lethal for fetuses preventing neonatal morbidity studies (Migale *et al*., 2015).

Recently, another study characterised a similar model of *E. coli*-induced PTB demonstrating bacterial ascension, maternal inflammation and early delivery, validating our results. However, regardless of *E. coli* dose, there were no surviving pups (Spencer *et al*., 2021). Others have modelled ascending infection with bacteria of particular relevance to human PTB such as *Ureaplasma parvum* and *Group B Streptococcus* (Pavlidis et al., 2020; Vornhagen et al., 2018; Bernardini et al., 2017; Tantengco et al., 2022; Randis et al., 2014). However, these bacteria do not consistently induce labour in mice and the impact on pups has not been investigated beyond recording survival immediately after birth. Although *E. coli* is not a common organism implicated in PTB, it is associated with preterm sequelae in mothers and is often associated with poor neonatal outcomes (McDonald *et al*., 1991; Mendz, Kaakoush and Quinlivan, 2013; Min *et al*., 2021). Specifically, *E. coli* K1 strains are commonly acquired from the mother at birth and are associated with neonatal sepsis and meningitis (McCarthy *et al*., 2016).

### Neuropathology

Markers of neuroinflammation were evident during the perinatal period at both the mRNA and protein level in pups exposed to *E. coli*, which is in agreement with mouse models of intrauterine inflammation induced by LPS (Brown *et al*., 2019; Burd *et al*., 2010; Burd *et al*., 2011; Elovitz *et al*., 2011; Chang *et al*., 2011; Ginsberg *et al*., 2021; Nadeau-Vallée *et al*., 2017). The developing brain is extremely sensitive to inflammation; an adverse environment even in an asymptomatic mother could cause brain injury in the fetus and cytokines are known to be neurotoxic (McAdams and Juul, 2012). Maternal infection and inflammation may alter the permeability of the fetal blood brain barrier, leading to neuroinflammation. Through exposure to excess inflammatory mediators, immune cells can accumulate at the embryonic choroid plexus, weakening tight junctions at this barrier (Cui *et al*., 2020). Although we see a trend for reduced ramification of microglia at E18.5, suggesting phagocytic activity, neuropathology is not fully evident until P7, which is likely the result of this perinatal neuroinflammation.

Only minimal inflammation, restricted to the prefrontal cortex region of the brain, remained by P7 with modest increases in *Il-1b* and the microglial marker *Iba1*. Interestingly, all mRNA expression was consistently reduced in the mid brain region (consisting mainly of the hippocampus, thalamus and striatum) of these mice. Downregulated mRNA expression could be the result of substantial cell death; there were activated microglia in the hippocampus, evidenced by a phagocytic phenotype, which may be associated with the increased TUNEL+ cell death and reduced neuronal density in this region.

Astrogliosis can occur in response to neuronal loss following central nervous system (CNS) injury. These cells release cytokines that may exacerbate inflammation and injury but they also restore homeostasis and promote neurogenesis (Pekny and Pekna, 2014). We do not see an increase in GFAP staining in the hippocampus at P7, which could suggest that neurogenesis was not stimulated by astrocytes in this region resulting in the consistently reduced neuronal density we observe at P7 and P14. However, our understanding is limited by the crude separation of the brain for mRNA analyses.

In addition to reduced neurogenesis, we observed increased activation of both microglia and astroglia across multiple brain regions, which has been reported in intrauterine inflammation models (Hester *et al*., 2018; Dada *et al*., 2014; Lei *et al*., 2017). We also see TUNEL+ cell death throughout the brain. Chorioamnionitis-associated white matter injury is common in preterm infants and is thought to begin *in utero* (Anblagan *et al*., 2016). It is associated with neuronal cell death, microglial activation and disruption to oligodendrogenesis, which can result in a myelination deficit (Leviton and Gressens, 2007; Anblagan *et al*., 2016). In our model, we observed increased microglial activation, increased cell death and increased astrogliosis in the external capsule at P7, suggesting white matter injury pathognomonic of PTB brain injury.

By P14, there is no longer measurable cell death or astrogliosis but neuronal density is still reduced in the in the hippocampus and also in the cortex. This suggests that neuroinflammation may have improved but there is a lasting effect on neurogenesis. We did not observe a disruption to myelination at P14 but an intrauterine LPS mouse model did not report this change until P30 (Chang *et al*., 2011).

Cortical area was reduced in *E. coli* exposed pups, similarly to prematurely born humans where reduced cortical thickness has been described (Nagy, Lagercrantz and Hutton, 2011; Dimitrova *et al*., 2021; Schmitz-Koep *et al*., 2020). Neuroimaging has confirmed reduced white and grey matter volumes in preterm infants and the microstructure of the cerebral cortex is altered in regions associated with cognition and sensory and motor function (Bouyssi-Kobar *et al*., 2018; Boardman and Counsell, 2020). We did not see a reduction in hippocampal volume, which has been reported in other animal models (Dada *et al*., 2014; Strahle *et al*., 2019).

Here, we have shown that exposure to ascending vaginal infection during gestation can cause disrupted neurodevelopment, which in humans can lead to cognitive and behavioural issues, with an increased the risk of cerebral palsy, attention deficit hyperactivity disorder, autism spectrum disorder and psychiatric issues (Johnson and Marlow, 2017).

### Lung injury

Fetal lungs are susceptible to injury when exposed to infection and/or inflammation via the amniotic fluid (Jackson *et al*., 2020). This results in an influx of neutrophils, increased inflammation and can impact alveolarization and surfactant production (Wirbelauer *et al*., 2008). Interestingly, inflammatory mediator mRNA expression was only upregulated in specific pups in our model. This could be a result of the positioning of the pups within the uterine horns. Nevertheless, we did observe elevation of certain secreted inflammatory mediators and the average number of neutrophils in the lungs was increased, which suggests that a measurable level of inflammation was occurring. Indeed, other models have reported an increase in inflammatory mediators (Yavuz *et al*., 2020; Papagianis *et al*., 2021; Kallapur *et al*., 2013) and neutrophil influx into the fetal lungs (Hudalla *et al*., 2018; Kallapur *et al*., 2013).

Pulmonary surfactants, which reduce surface tension, are essential for respiration (Meyer and Zimmerman, 2002). We observed reduced surfactant protein mRNA expression in the perinatal pup lungs, which is supported by data from other murine models (Yavuz *et al*., 2020; Salminen *et al*., 2008). Surfactant deficiency and lung morphological disruption can result in respiratory distress syndrome (RDS) and consequent ventilatory support, which can subsequently increase susceptibility to conditions such as BPD and chronic lung disease of prematurity (Davidson and Berkelhamer, 2017). We observed morphology disruption in the lungs evidenced by increased alveolar area, which has been reported in mouse, sheep and non-human primate models (Habelrih *et al*., 2022; Papagianis *et al*., 2021; Toth *et al*., 2022), which may be the result of activated immune cells in the lung (Stouch *et al*., 2016).

A limitation of using mice to model prenatal lung injury is that alveologenesis occurs postnatally in mice rather than before term as in humans and non-human primates (Jackson *et al*., 2020). This could explain why we don’t see accelerated lung maturation; nevertheless, our model does appear to recapitulate the morphological changes seen in neonatal BPD models (Durrani-Kolarik *et al*., 2017; Nguyen *et al*., 2019).

### Gastrointestinal inflammation

We observed a marked increase in inflammatory cytokine mRNA expression in the gut. This particular strain of *E. coli* is known to colonise the gastrointestinal tract, which has been demonstrated in neonatal rats. The bacteria cross the epithelium of the gut into the systemic circulation and enter the CNS by crossing the blood brain barrier, via choroid plexus epithelium, where they invade the meninges (McCarthy *et al*., 2016; Zelmer *et al*., 2008; Witcomb *et al*., 2015). One study demonstrated increased gut inflammation and increased gut permeability in naturally preterm pups, so even in the absence of infection prematurity alone is enough to disrupt the gut (Arboleya *et al*., 2021).

Comparatively, a neonatal mouse model of NEC describes disruption to the intestinal mucosa and increased *Il-1β* and *Cxcl2* expression (Mihi *et al*., 2021). As the immune system of premature babies may be underdeveloped or dysregulated, they are susceptible to conditions such as NEC resulting from an exacerbated inflammatory response to microbial products introduced during feeding (Henderickx *et al*., 2019). In addition, premature infants have delayed bacterial colonisation and reduced microbial diversity in the gut, which can be affected up to four years following disruption during the perinatal period (Rougé *et al*., 2010; Arboleya *et al*., 2012; Fouhy *et al*., 2019). NEC, itself, can result in brain damage due to increased permeability of the gut caused by damage to the intestinal epithelium. Bacteria can translocate into the blood, which can lead to brain damage. This could be due to the activation of T cells, neurotoxic levels of IFNγ, which can activate microglia and lead to disrupted myelination (Zhou *et al*., 2021). Although NEC is believed to be established postnatally, we may be modelling a similar pathology. In addition, the colonisation of the gut postnatally may contribute to the neonatal neuropathology we observe.

### Validating the model by cervical HBD3 gene therapy

We hypothesised that by facilitating the upregulation of the antibacterial peptide HBD3, we would enhance the immunological barrier of the cervix, thus protecting the developing fetus from morbidity and mortality. Using our model of ascending vaginal infection, we have validated our previous finding that this gene therapy increases the number of live born pups (Suff *et al*., 2020). We have expanded on these results, confirming that postnatal pup survival is also improved and there is a reduction in *Tnfα* mRNA expression in the perinatal pup brain, suggesting that this treatment may shield pups from the damaging levels of inflammation resulting from maternal infection. Human neonatal brain damage is associated with elevated TNFα in cord blood (Wikström *et al*., 2008; Lu *et al*., 2016; Lee *et al*., 2021). Activated microglia release TNFα but these cells can also be activated by TNFα via a positive feedback loop, resulting in sustained activation (Kuno *et al*., 2005). Therefore, by reducing *Tnfα* this treatment may prevent neuropathology mediated by cytokine toxicity and microgliosis. Although HBD3 gene therapy did not prevent PTB in this model, it played an arguably more important role in improving pup survival and reducing inflammation in the perinatal brain and, therefore, this treatment strategy warrants further investigation.

### Conclusion

It is essential to develop clinically relevant animal models, such as the one we have described, in order to investigate the underlying mechanisms leading to PTB and for the advancement of novel therapies. Here, we demonstrate that our mouse model of ascending vaginal infection induces PTB and reduces pup survival but, importantly, this model also recapitulates neonatal morbidity. This includes lung injury, gut inflammation and neonatal neuropathology; all of which are fundamental to the conditions premature babies are susceptible to. We also demonstrated how this model can be used to investigate novel therapeutics targeting PTB and the associated neonatal morbidities.

## Author contributions

AB: conception and design, acquisition, analysis and interpretation of data, drafted and revised the manuscript. KT: acquisition, analysis and interpretation of data, manuscript revision. NS: conception and design, acquisition, analysis and interpretation of data, manuscript revision. LB: acquisition of data. MM: acquisition of data. MH: experimental design and interpretation of data. SW: conception and design, analysis and interpretation of data, manuscript revision. DP: conception and design, interpretation of data, manuscript revision.

## Acknowledgments

This project was supported by Action Medical Research and The Borne Foundation (GN2647) and Wellbeing of Women (RG2365).

## Figure legends

**Supplementary figure 1. Perinatal neuropathology assessments.** Pyriform cortex (Pyr), external capsule (EC), striatum (Str), hippocampus (Hip), corpus callosum (CC), thalamus (thal).

**Supplementary figure 2. M1 and M2 assessments of the perinatal pup brain.** M1 is reduced in the thalamus of the *E. coli* group (A). Pyriform cortex (Pyr), striatum (Str), hippocampus (Hip), corpus callosum (CC), thalamus (thal).

**Supplementary figure 3. P14 neuropathology assessments.** Pyriform cortex (Pyr), external capsule (EC), striatum (Str), hippocampus (Hip), thalamus (thal), total (all regions), optical luminosity values (OLV).

**Supplementary figure 4. Neonatal lung assessments.** We did not observe changes to the lung morphology or neutrophil influx at P7.

