## Supplementary figures and images for "Ascending vaginal infection in mice induces preterm birth and neonatal morbidity"

### Supplemental figures

#### Supplementary figure 1

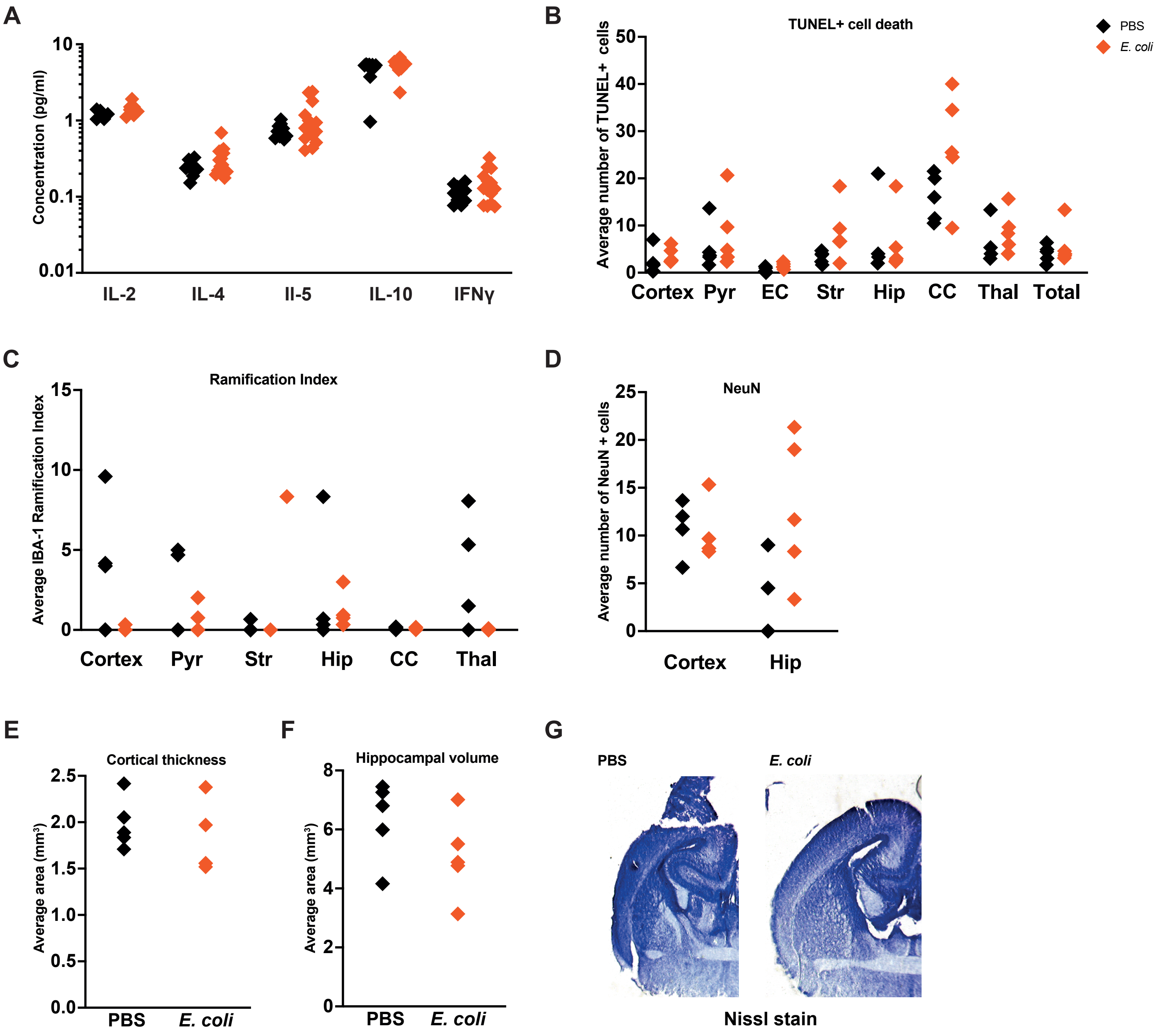

#### Supplementary Figure 2

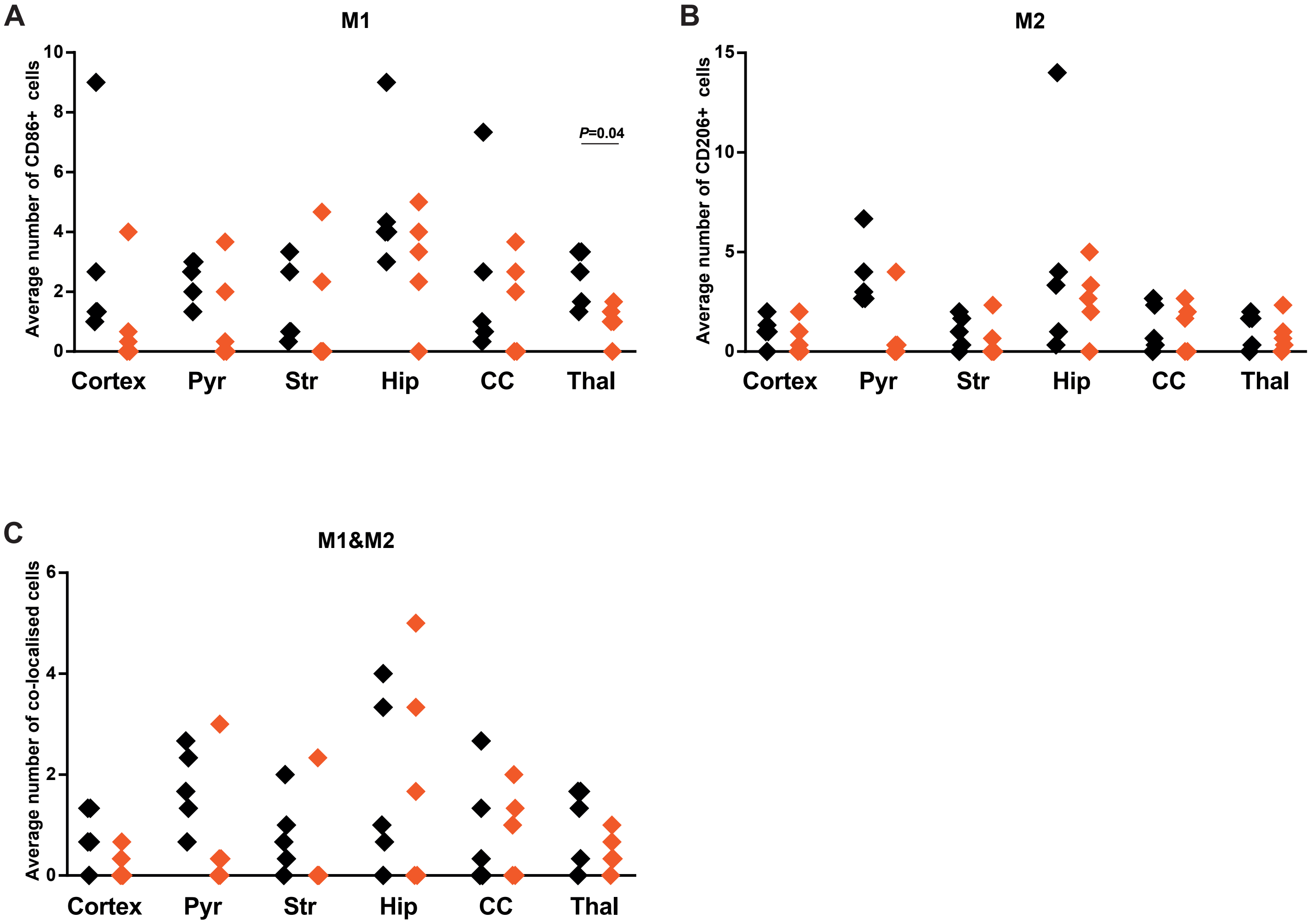

#### Supplementary Figure 3

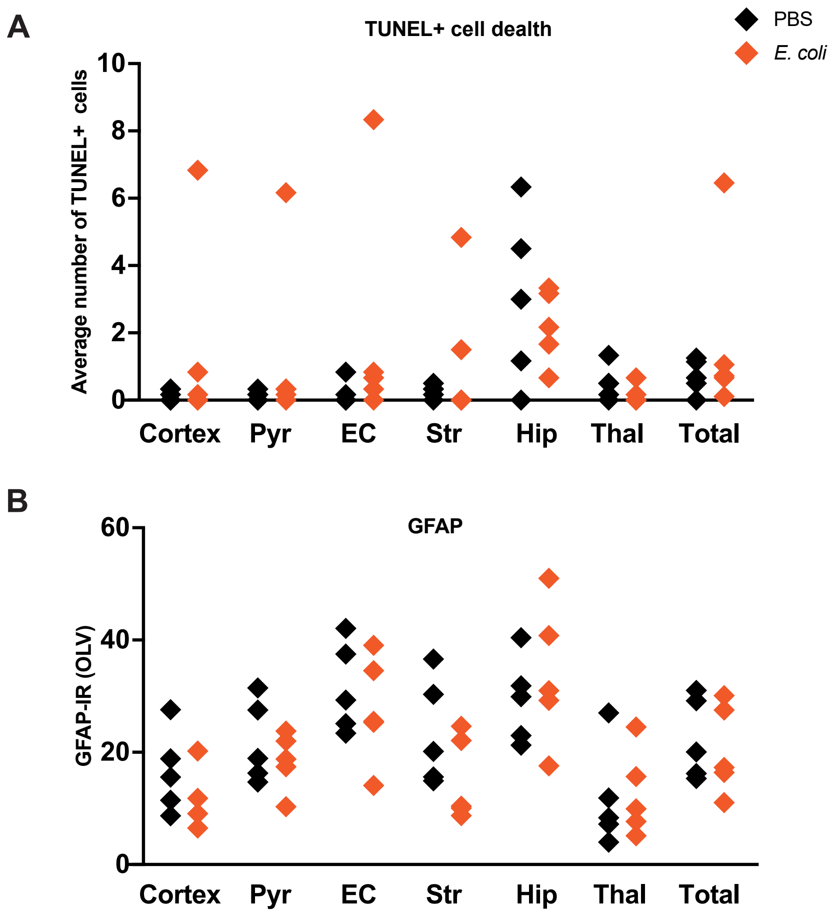

#### Supplementary Figure 4

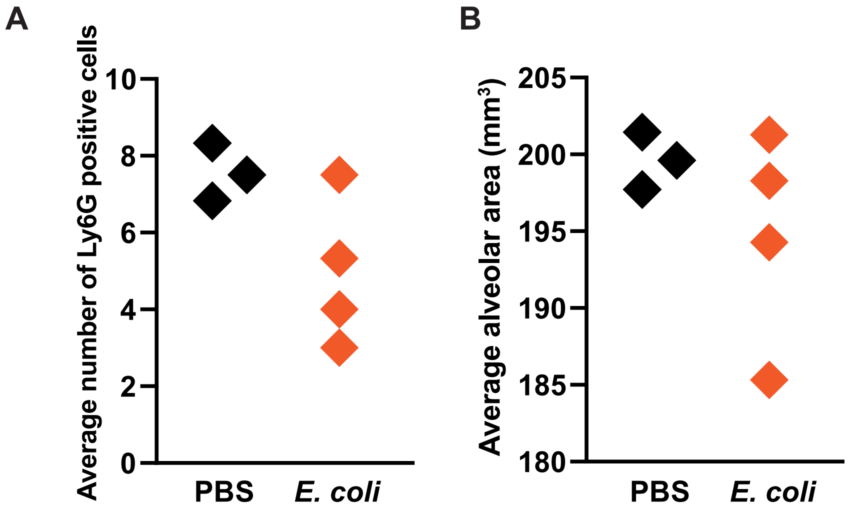
